# ALiCE^®^: A versatile, high yielding and scalable eukaryotic cell-free protein synthesis (CFPS) system

**DOI:** 10.1101/2022.11.10.515920

**Authors:** Mainak Das Gupta, Yannick Flaskamp, Robin Roentgen, Hannes Juergens, Jorge Armero Gimenez, Frank Albrecht, Johannes Hemmerich, Zulfaquar Ahmad Arfi, Jakob Neuser, Holger Spiegel, Alexei Yeliseev, Lusheng Song, Ji Qiu, Charles Williams, Ricarda Finnern

**Author notes:** Corresponding author: Dr. Ricarda Finnern.

## Abstract

Eukaryotic cell-free protein synthesis (CFPS) systems have the potential to simplify and speed up the expression and high-throughput analysis of complex proteins with functionally relevant post-translational modifications (PTMs). However, low yields and the inability to scale such systems have so far prevented their widespread adoption in protein research and manufacturing.

Here, we present a detailed demonstration for the capabilities of a CFPS system derived from *Nicotiana tabacum* BY-2 cell culture (BY-2 lysate; BYL). BYL is able to express diverse, functional proteins at high yields in under 48 hours, complete with native disulfide bonds and N-glycosylation. An optimised version of the technology is commercialised as ‘ALiCE^®^’, engineered for high yields of up to 3 mg/mL. Recent advances in the scaling of BYL production methodologies have allowed scaling of the CFPS reaction. We show simple, linear scale-up of batch mode reporter proten expression from a 100 μL microtiter plate format to 10 mL and 100 mL volumes in standard Erlenmeyer flasks, culminating in preliminary data from 1 L reactions in a CELL-tainer® CT20 rocking motion bioreactor. As such, these works represent the first published example of a eukaryotic CFPS reaction scaled past the 10 mL level by several orders of magnitude.

We show the ability of BYL to produce the simple reporter protein eYFP and large, multimeric virus-like particles directly in the cytosolic fraction. Complex proteins are processed using the native microsomes of BYL and functional expression of multiple classes of complex, difficult-to-express proteins is demonstrated, specifically: a dimeric, glycoprotein enzyme, glucose oxidase; the monoclonal antibody adalimumab; the SARS-Cov-2 receptor-binding domain; human epidermal growth factor; and a G protein-coupled receptor membrane protein, cannabinoid receptor type 2. Functional binding and activity are shown using a combination of surface plasmon resonance techniques, a serology-based ELISA method and a G protein activation assay. Finally, in-depth post-translational modification (PTM) characterisation of purified proteins through disulfide bond and N-glycan analysis is also revealed - previously difficult in the eukaryotic CFPS space due to limitations in reaction volumes and yields.

Taken together, BYL provides a real opportunity for screening of complex proteins at the microscale with subsequent amplification to manufacturing-ready levels using off-the-shelf protocols. This end-to-end platform suggests the potential to significantly reduce cost and the time-to-market for high value proteins and biologics.

## Introduction

Proteins are critical components of medicines, vaccines and diagnostics. However, the design-build-test-learn cycles are too slow for protein research and manufacture using current cell-based methods. As highlighted by the recent COVID-19 pandemic, there is a need for end-to-end platforms that enable rapid research and development (R&D) pipelines with subsequently scalable manufacturing processes, to enable society to react rapidly to future threats to public health.

Cell-free protein synthesis (CFPS) has emerged as a powerful approach with the potential to enable accelerated R&D workflows, by first providing rapid and simple protein screening and later, the scaled production of promising protein leads ^1–3^. In comparison to cell-based expression, CFPS offers unprecedented speed with weeks of cell culturing, engineering, clone selection and expansion, condensed into hours ^1,4^. Typically comprising a cell lysate or extract, a catalogue of CFPS platforms have been developed in the past decades and are defined by their starting cellular material. These include systems derived from bacterial, yeast, mammalian and insect cell cultures, Wheat Germ Extract (WGE) and reticulocytes from rabbit blood^5^.

By decoupling the concerns of biomass accumulation and cell culture constraints from the essential mechanisms of recombinant protein production, CFPS offer unique advantages, such as the cytotoxic over-expression of membrane proteins and peptides which are often tightly regulated within intact cells^6–8^. However, the widespread adoption of CFPS in therapeutic discovery pipelines has not materialized, due to: low product yields, inadequacies with installing post-translational modifications (PTMs) that are essential for correct folding and function of complex proteins, and a lack of scalability in cell lysate production methodologies and corresponding CFPS reactions.

A eukaryotic, coupled transcription-translation CFPS system based on tobacco BY-2 cell lysate (BYL) promises to overcome these issues. Following initial publications^9,10^, BYL has since been optimised and commercialised by LenioBio GmbH as ALiCE^®^ (Almost Living Cell-free Expression). As BYL contains native, actively-translocating microsome vesicles derived from the endoplasmic reticulum and Golgi, it outperforms other CFPS modes in its capability of performing PTMs^9,10^. Reporter protein yields are also notably higher than those reported for other systems^5^. The use of relatively inexpensive plant cell culture as the feedstock for lysate production combines with a relatively simple bioprocessing method to allow much better scaling potential compared to closely related eukaryotic CFPS systems e.g. WGE produced from full plants. At the same time, BYL reactions are high-yielding in batch mode, bypassing complex and expensive continuous-exchange cell-free formats that are needed to make other eukaryotic CFPS reactions viable^11–13^.

Indeed, recent advances in BYL manufacturing have allowed for scaling of the BYL CFPS reaction. Here, we show for the first time how a eukaryotic CFPS reaction can easily and linearly scale across a 1000x fold volume difference from 100 μL reactions to 10 mL and 100 mL, for both the cytosolic reporter eYFP and the multi-domain microsomal glycoprotein glucose oxidase (GOx). Preliminary data from 1 L reactions are also disclosed, representing the first such example of litre scale reactions from a eukaryotic CFPS. No significant difference in yield was observed in direct comparison with 50 μL reactions, highly remarkable considering the 20,000x fold volume difference

Scaling of protein reactions consequently enables the purification of useful amounts of sample for more in-depth characterisation of the proteins produced in BYL. Thus, we report the expression and characterization of an array of difficult-to-express proteins, specifically: GOx; a hepatitis B core virus-like particle (HBc-VLP), the monoclonal antibody adalimumab; the SARS-CoV-2 receptor-binding domain (RBD); human epidermal growth factor (hEGF),and a G protein-coupled receptor membrane protein, cannabinoid receptor type 2 (CB2) (Figure 1). This panel features proteins of monomeric and multimeric nature, with disulfide bonds, N-glycosylation and transmembrane domains, thus displaying the versatility of BYL. Importantly, we also present mass spectrometry analysis of disulfide bond formation and N-glycan structural characterisation for some of these proteins. Such detailed analysis has never before been shown for proteins from eukaryotic CFPS, due to previous limitations in reaction volumes and yields. Here we found correct disulfide bond formation and high occupancy, homogeneous N-glycan profiles with high mannose structures typical of plant-derived recombinant proteins. Finally, these data are complemented by functional assays, to include surface plasmon resonance (SPR) binding analyses of adalimumab and hEGF against their cognate receptors, a serology-based ELISA of RBD against SARS-CoV-2 patient samples and a G protein activation assay using CB2 directly in the lysate matrix.

**Figure 1.**
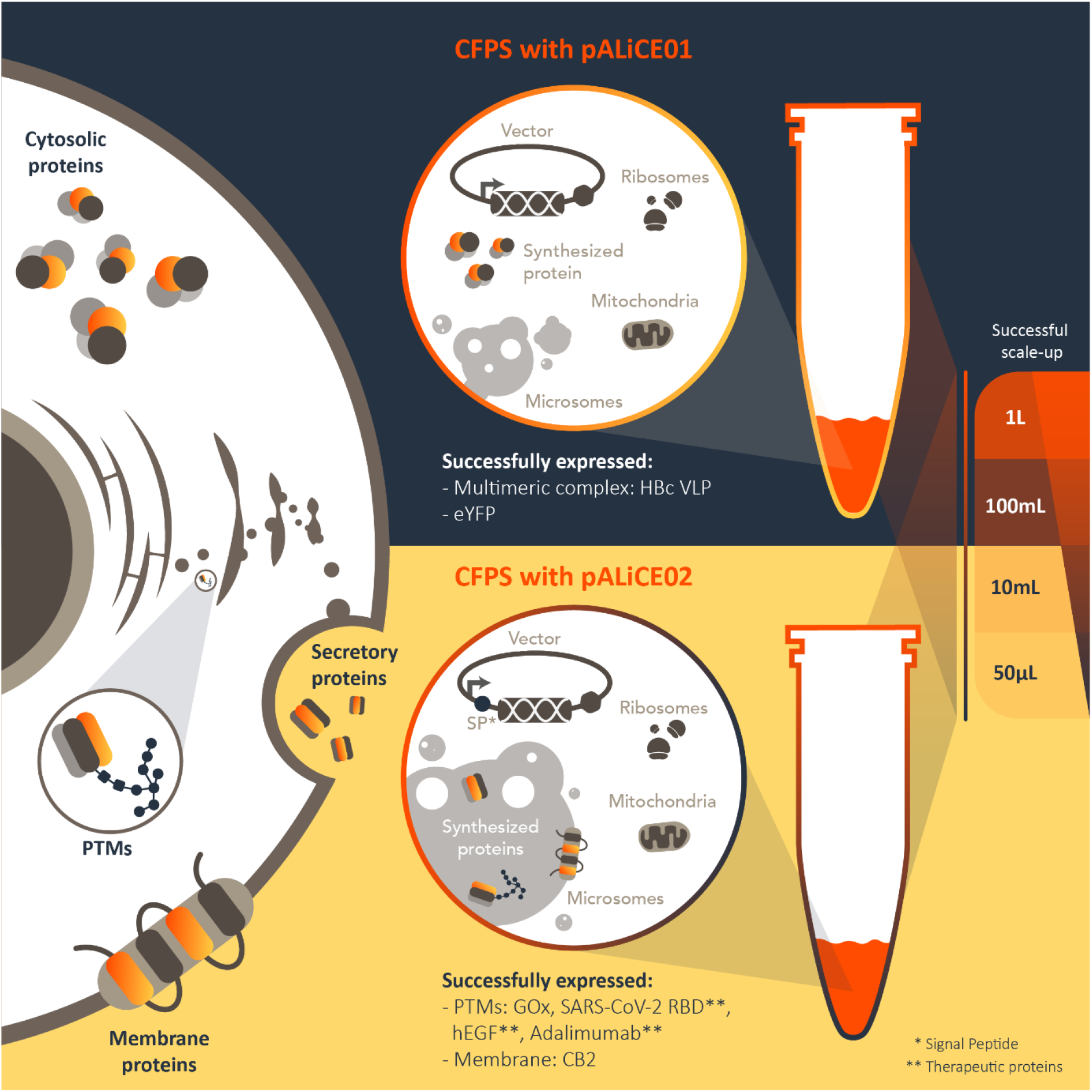
Cytosolic and microsomal protein expression in ALiCE. Cytosolic expressed proteins (top) include Hepatitis B core antigen virus-like particle (HBc VLP) and Enhanced Yellow Fluorescent Protein (eYFP). Microsomal expressed proteins (bottom), include Glucose Oxidase (GOx), the receptor binding domain of the SARS-CoV-2 spike protein (RBD), human Epidermal Growth Factors (hEGF), the antibody, Adalimumab, and the multi-pass transmembrane G protein-coupled receptor, cannabinoid receptor 2 (CB2).

## Results

### Linear Scalability of CFPS Reactions in BYL

BY-2 cell lysate (BYL) is commercialised as the product ALiCE®, optimised for the capability to produce 3 mg/mL of the cytosolic reporter protein eYFP in sub-millilitre volumes (Figure 2A). This yield in small volumes is ideal for screening and research purposes. However, significantly larger reaction volumes are necessary to produce proteins at the amounts required for industrial applications. Scaling of CFPS systems first requires scaling of the cellular input material fermentation volumes and of the lysate production process, in order to generate sufficient lysate material to allow for scaling of the actual CFPS reaction. This reaction scaling requires an exploration of reaction vessels, oxygen transfer rates and other protein expression reaction conditions^5^.

**Figure 2.**
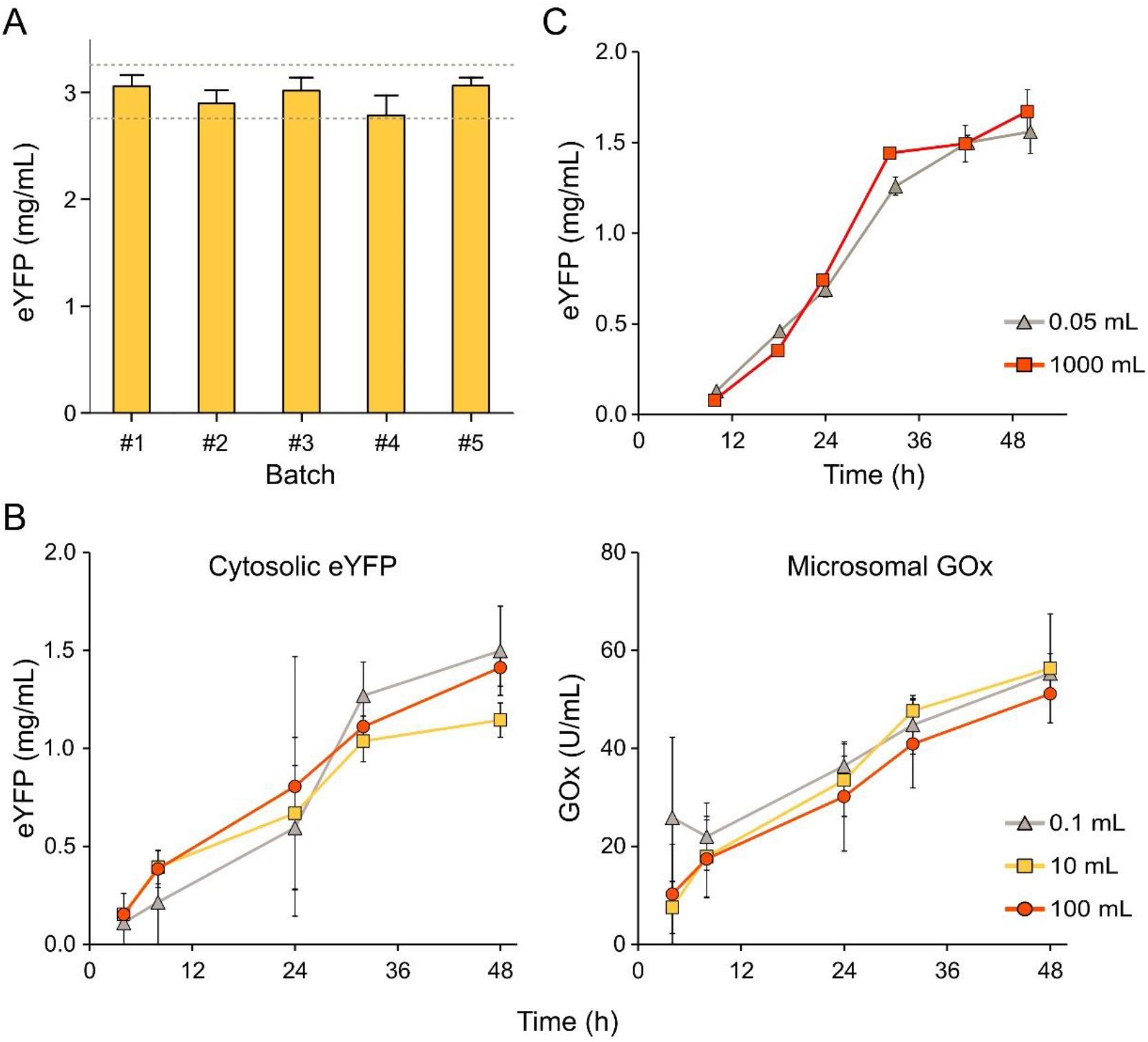
Scaling cell-free protein synthesis reactions using BY-2 lysate. A) eYFP expression yields in recent commercial batches of ALiCE^®^ with release criteria of 2.7-3.3 mg/mL shown. Fluorescence of lysate samples are compared against eYFP standards to calculate yields. B) Cytosolic and microsomal protein expression scaling at 0.1 mL, 10 mL and 100 mL. CFPS reactions were performed in either microtiter plates or shake flasks for 48 hours and at 25 °C, with eYFP and GOx concentrations determined by fluorescence and colorimetric activity assays, respectively (N=2). C) Preliminary trial of a 1000 mL CFPS reaction for cytosolic eYFP production. Comparison with microplate reactions of 50 μL show no significant difference between protein yield across a scaling factor of 20,000x.

Scaled BY-2 culture and lysate production has been achieved and now permits liter scale BYL production. Thus, we were able to test the consistency of BYL performance across reaction volumes of 0.1, 10 and 100 mL using just two large batches of lysate for all scaling experiments. pALiCE01 and pALiCE02 vectors encoding the reporter proteins eYFP and GOx, respectively, were used as template DNA material for 48 hour cytosolic and microsomal CFPS reactions at these different scales (Figure 2B). eYFP and GOx titers demonstrated strikingly linear scalability from 0.1 to 100 mL reaction volumes, demonstrating a scaling factor of 1000x with no significant loss of protein yield at the reaction endpoint, regardless of expression mode and protein. These data represent the largest eukaryotic CFPS reactions to date and the linear scaling supports the use of BYL as an end-to-end R&D platform, whereby protein engineering can be performed at the microliter scale before undemanding upscaling.

For 0.1 mL expressions, microtiter plates were used, whilst orbitally shaken Erlenmeyer-type glass flasks were chosen as milliliter reaction vessels due to the universality of this glassware and the availability of characterisation data regarding oxygen transfer rate (OTR) and global power input^14,15^. Considering no difference in scalability potential between the simple, cytosolic eYFP and the complex, microsomal GOx, it is not expected that these simple conditions need further optimisation when moving across the micro-milliliter scale. However, to truly achieve manufacturing potential for high value proteins, eukaryotic CFPS reactions must also be scaled to liters and potentially kiloliters in the future. Thus, we also present the results of a preliminary trial comparing 50 μL microplate reactions to a 1 L BYL reaction using the commercial bioreactor, the CELL-tainer® CT20, using the same batch of lysate at both scales (Figure 2C). Remarkably, timecourse data reveals that eYFP yields were practically identical throughout the 48 hour runtime, regardless of reaction scale. eYFP yields for these reactions peaked at 1.5 mg/mL, suggesting that the scaled processing methodology needs optimisation to recoup the 3 mg/mL potential of BYL. Nonetheless, the consistency of yield across a scaling factor of 20,000x is a clear indication of the potential for this eukaryotic CFPS system.

### Expression of multisubunit protein complexes in BYL with in-depth molecular and functional characterisation

Virus-like particles (VLPs) are complex nanostructures built by the specific interaction between hundreds of virus-derived protein monomers. As they closely resemble viruses but are unable to infect the host cell due to absence of viral genetic material, VLPs can be utilized in medicine as potent and safe vaccines^16^. Furthermore, VLPs may serve as delivery vectors for RNA/DNA vaccines or targeted drug delivery^17^. Production of VLPs by CFPS has the added advantage that reaction parameters can be readily modified to improve VLP assembly rates. However, previously published efforts with eukaryotic CFPS have struggled to achieve useful yields in meaningful reaction volumes. In the case of the Hepatitis B core antigen model VLP (HBc), *Pichia pastoris* and wheat-germ extract (WGE) systems have yielded only 6.4 and 4 μg/mL VLP production, respectively, and neither system was used beyond the microliter scale for HBc^18–20^.

Using the BYL CFPS system, high concentrations of fully assembled HBc VLP were obtained after a 48h reaction, without the need for further optimization. SDS-PAGE analysis first confirmed the presence of the 21 kDa monomer in the lysate and after purification (Figure 3A). In-gel quantification gave an estimated HBc monomer yield of 1 mg/mL, representing 150-fold increases over other eukaryotic CFPS reports. Electron micrographs confirmed the presence of a chomogenous population of fully-formed VLPs in the lysate (Figure 3B). The BYL also outperformed HBc yields of 0.4 mg/mL from an *Escherichia coli* CFPS system^21^. These results show the potential of BYL to produce high-yields of multimeric complexes like VLPs.

**Figure 3.**
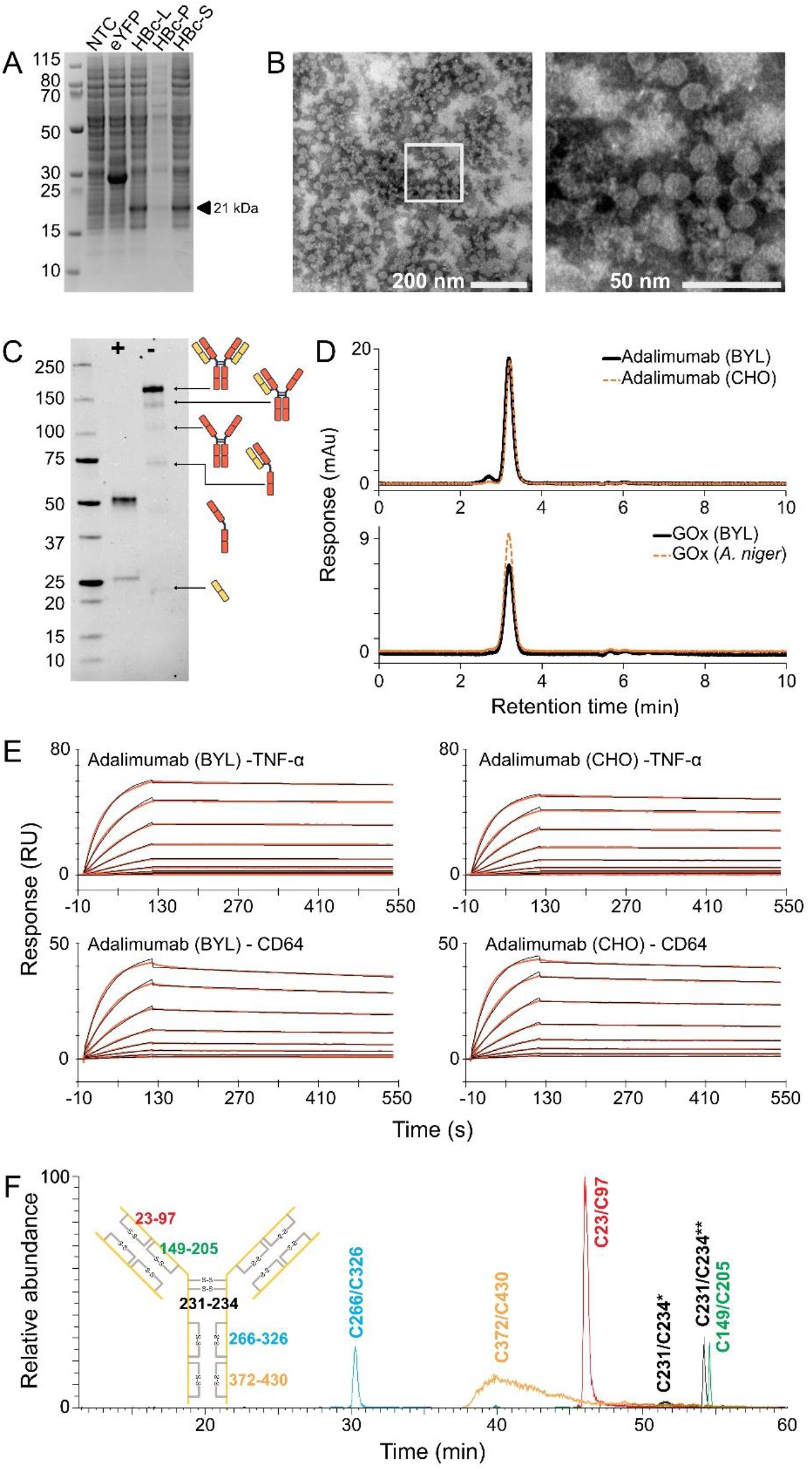
Production, characterisation and functional assessment of multisubunit complexes in BY-2 lysate (BYL). A) SDS-PAGE of Hepatitis B core antigen (HBc) monomer produced in BYL. The expected 21 kDa band is indicated by the arrow. NTC – non-template control; eYFP – eYFP control reaction; HBc-L – total lysate after expression; HBc-P – insoluble fraction pellet; HBc-S – soluble protein fraction; HBc – purified HBc sample. B) Electron microscopy picture of negatively stained, self-assembling virus-like particles in BYL. C) Anti-IgG Western blot of pure adalimumab electrophoresed under reducing (+) and non-reducing (-) conditions. D) SEC-HPLC analysis of adalimumab and GOx produced in BYL compared to commercial standards. E) SPR analysis of adalimumab and its binding partners. Immobilised antibody molecules produced from either BYL or CHO cells were probed with recombinant TNFa and CD64 to show ligand and receptor binding, respectively. F) Disulfide bond analysis of adalimumab.

Additional multisubunit model proteins were selected for further study of folding and post-translational modification capabilities, namely: the enzyme GOx, a homodimer comprised of 80 kDa monomers that are covalently linked by disulfide bonds and each possessed of 8 N-glycosylation sites; and the therapeutic monoclonal antibody adalimumab (Humira^®^), a complex heterotetramer of 150 kDa comprised of two heavy and two light chains, together containing two N-glycosylation sites, twelve intramolecular and four intermolecular disulfide bonds. Adalimumab heavy and light chain genes were cloned directly into pALiCE02 vectors for microsomal targeting whilst RBD and GOx genes were prepared similarly with additional N-terminal Strep-II tags for purification. Proteins were expressed in 10 mL reactions using triplicate, independent batches of BYL and the resulting microsomes were processed by dodecylmaltoside detergent treatment for protein release and subsequent Protein A or streptavidin affinity purification.

For adalimumab, a 1:1 ratio of heavy and light chain pALiCE02 plasmids was sufficient to produce full-length IgG molecules at the expected 150 kDa mass (Figure 3C). Analytical size-exclusion chromatography resolved a single major peak for Adalimumbab expressed in ALiCE indicating a homogeneous population of heterotetrameric mAb (Figure 3D). Indeed, a near-perfect overlay was observed between the chromatograms of Adalimumab produced cell-free in ALiCE as well as in CHO cells. Intermediate assemblies of heavy and light chain visible in the non-reducing PAGE (Figure 3C) were probably caused by partial dilsuphide bond disruption during protein migration through the gel matrix. Similarly comparable SEC profiles with single major peaks were seen for GOx expressed in BYL and a commercial standard purified from *Aspergillus niger*, indicating homogeneous dimeric protein molecule with no misprocessed variants or monomeric species observed for either sample (Figure 3D).

The functional binding activity of adalimumab samples towards the tumor necrosis factor (TNFα) ligand and the Fc-gamma receptor CD64 was assessed by SPR in a comparison with a commercial adalimumab reference product from CHO cells (Figure 3E). For TNFα binding, BYL-mAb demonstrated slightly stronger affinity than CHO-mAb, with an average K_D_ (M) value of 1.34×10^-10^ compared to 1.85×10^-10^, respectively. Conversely, BYL-mAb showed weaker binding to the CD64 receptor than CHO-mAb (3.53×10^-10^ compared to 1.58×10^-10^). Considering that the adalimumab mechanism of action is the neutralisation of free TNFα, this enhanced ligand binding and weaker receptor binding may combine for an improved therapeutic effect when this monoclonal antibody is produced in BYL compared to mammalian cell systems. Disulfide bond analysis of adalimumab was also used to demonstrate the correct formation of the expected intra- and intermolecular bonds (Figure 3F).

### N-linked glycosylation of glycoproteins expressed in BYL

To assess the N-glycosylation potential of the eukaryotic BYL a panel of glycoproteins was used, to include adalimumab and GOx, together with the receptor binding domain (RBD) of the SARS-CoV-2 spike, a monomer of 52 kDa with four intramolecular disulfide bonds and two N-glycosylation sites (Figure 4). Similar to previous reports of glycoproteins produced by eukaryotic CFPS^22^, the presence of N-glycosylation is demonstrated using simple SDS-PAGE mobility shifts after glycosidase treatment with either Peptide:N-glycosidase F or Endoglycosidase H (PNGase F and EndoH, respectively; Figure 4A). However, there has been no detailed analysis on the composition of these N-glycans as the requisite sample amounts for mass spectrometric characterisation were unobtainable. Successful scaling of the BYL system has enabled sufficient sample amounts for more in-depth examination and so we analysed the N-glycan structures of the GOx, RBD and adalimumab preparations that were also used for disulfide bond characterisation.

**Figure 4.**
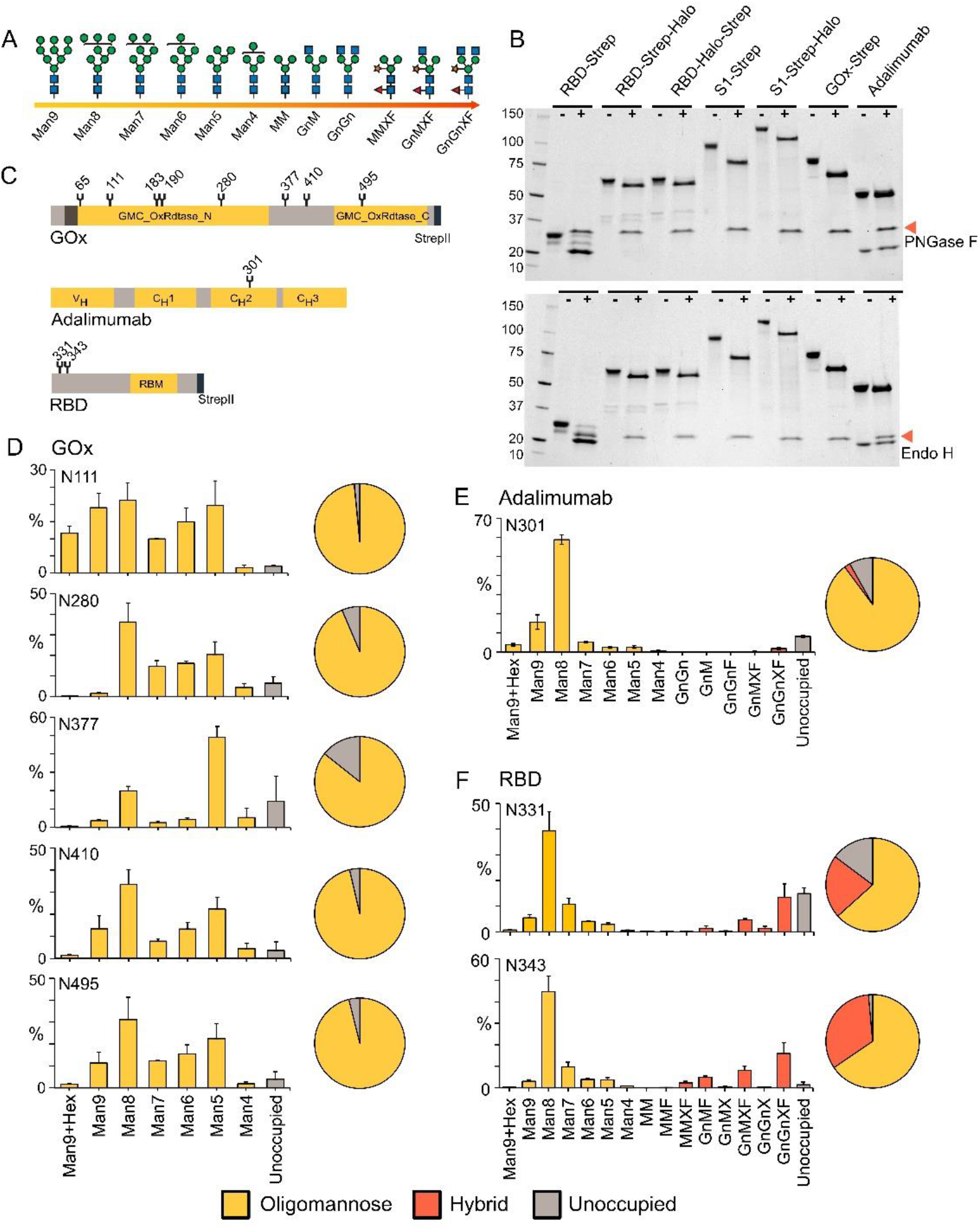
N-glycosylation investigation of proteins produced in BYL. A) Schematic explanation of N-glycan structures and nomenclature. B) SDS-PAGE mobility shift after glycosidase treatment. PNGase F – Peptide:N-glycosidase F; EndoH – Endoglycosidase H. C) Schematic representation of protein N-glycosites. D) Summary of LC-ESI-MS/MS analysis of N-glycan structures for adalimumab, SARS-CoV-2 receptor binding domain (RBD) and GOx glycosites.

N-glycopeptide digestions were followed by liquid chromatography electrospray ionization tandem mass spectrometric identification and assignment of structures (LC-ESI-MS/MS; Figure 4D). The analysis revealed a high degree of N-glycosylation occupancy and reproducibility across batches, regardless of the protein model. The N-glycan profile is broadly consistent across all N-glycosites and demonstrates a dominant mannose 8 (Man8) species. Similarly to recombinant proteins produced in whole *Nicotiana* spp. plants, plant-specific glycosylation features of α1, 3-fucosylation and β1, 2-xylosylation were also observed at some N-glycosites, especially within the RBD protein. One glycosylation position on the GOx protein was not detected and, therefore, not characterized. This site was located within a weakly charged peptide of approximately 50 amino acids, presumed to have hindered detection within the mass range of the MS instrument.

### Functional expression of a multi-pass transmembrane G protein-coupled receptor, cannabinoid receptor 2 (CB2)

Integral membrane proteins are challenging for recombinant expression due to a number of specific requirements for folding, PTMs, and insertion of these proteins into cell membranes or detergent micelles^23,24^. G protein-coupled receptors (GPCRs) are one of the largest classes of integral membrane proteins involved in a wide array of signal transduction and regulatory pathways in the human body, and therefore attract significant attention. However, despite significant efforts over the past two decades, there has been limited success in expressing correctly folded and functionally active GPCR in CFPS systems^25^. An improved methodology has been described for functional *E. coli* expression of the class A rhodopsin-like GPCR human cannabinoid receptor type II (CB2), as well as from mammalian HEK293 and CHO cell cultures^26–28^. However, no successful CFPS of the CB2 receptor has been reported until now.

Figure X presents proof of functional CB2 expression in BYL. The 44 kDa CB2 was expressed from both pALiCE01 and pALiCE02 vectors as a fusion with the N-terminal maltose-binding protein (MBP) of *E. coli*, a Strep-II tag and a C-terminal His-tag. CB2 expression levels were measured by Western blot and function was demonstrated by activity assays. The Western blot image (Figure 5A) shows BYL reaction products, probed with anti-CB2 (left panel) and anti-Strep-II tag (right panel) antibodies. CB2 is well expressed from both pALiCE01 and pALiCE02 vectors, with estimated yields of 150-200 μg/mL.

**Figure 5.**
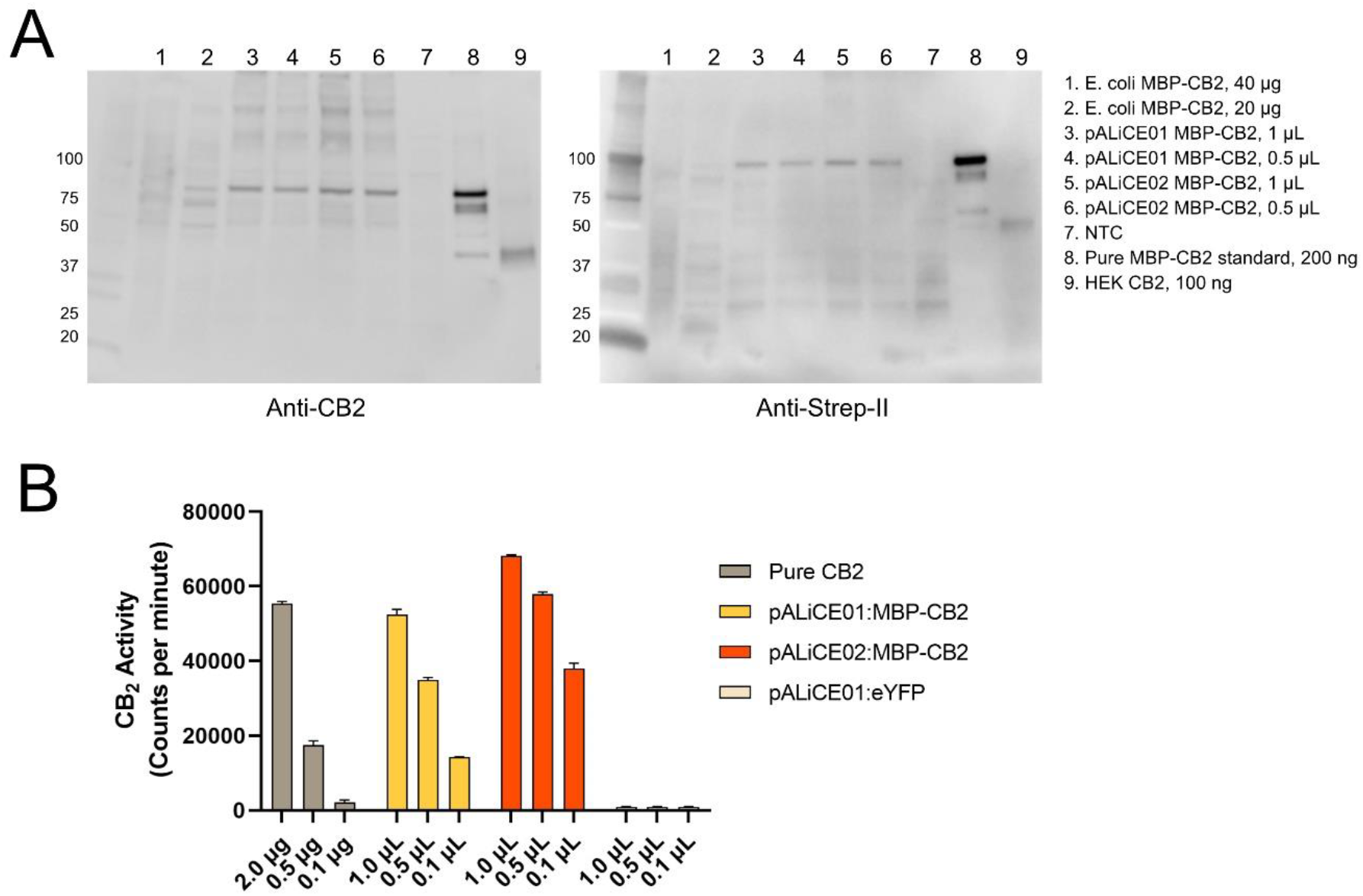
Expression and activity assays of human cannabinoid receptor CB2 expressed in BYL. A) Western blotting with anti-CB2 and anti-Strep-II tag antibodies. A DNA-free negative control and MBP-CB2 positive controls from *E. coli* expression are included as indicated. B) CB2 activation assay from CP-55,940 scintillation counts. Bars represent average values and standard deviations from four independent measurements.

Functional activity of the expressed CB2 protein was assessed in the lysate by measurement of G protein activation rates upon binding of the receptor with the synthetic cannabinoid agonist CP-55,940 (Figure 5B). Remarkably, functional protein was found from both pALiCE01 and pALiCE02 expression vectors, possibly suggesting a passive loading mechanism of the protein to microsomal membranes in the former case. Comparison with recombinant CB2 purified from *E. coli* gives approximate BYL yields of 150-200 μg/mL without any optimization of reaction conditions or the addition of supplementary lipids, detergents or nanodiscs that are required for other CFPS systems without native membrane compartments.

### Functional expression of therapeutic proteins – a SARS-CoV-2 vaccine subunit candidate and human epidermal growth factor

CFPS is well suited to the study of emerging pathogens, with rapid expression times that can outpace even quickly mutating threats. A concerted, global research effort was essential for mitigating the recent SARS-CoV-2 pandemic but a preponderance of expensive, low quality commercial antigen samples did not help. Antigens from mammalian cells, yeast cells and baculovirus-insect cells proved costly and too low yielding to support vaccine or therapeutic development, while antigens derived from heterologous *E. coli* expression were highly available but lack the post-translational modifications of the disease-relevant proteins^29^. Herein we have shown the BYL system to be capable of producing recombinant SARS-CoV-2 spike protein RBD within 48 hours and with consistent batch-to-batch N-glycosylation. The RBD was expressed in BYL and purified from BYL with a high degree of purity (Figure 6A). The sample that was used for N-glycopeptide analysis was also subject to disulfide bond analysis, again confirming that BYL produces proteins with the correct folding (Figure 6B).

**Figure 6.**
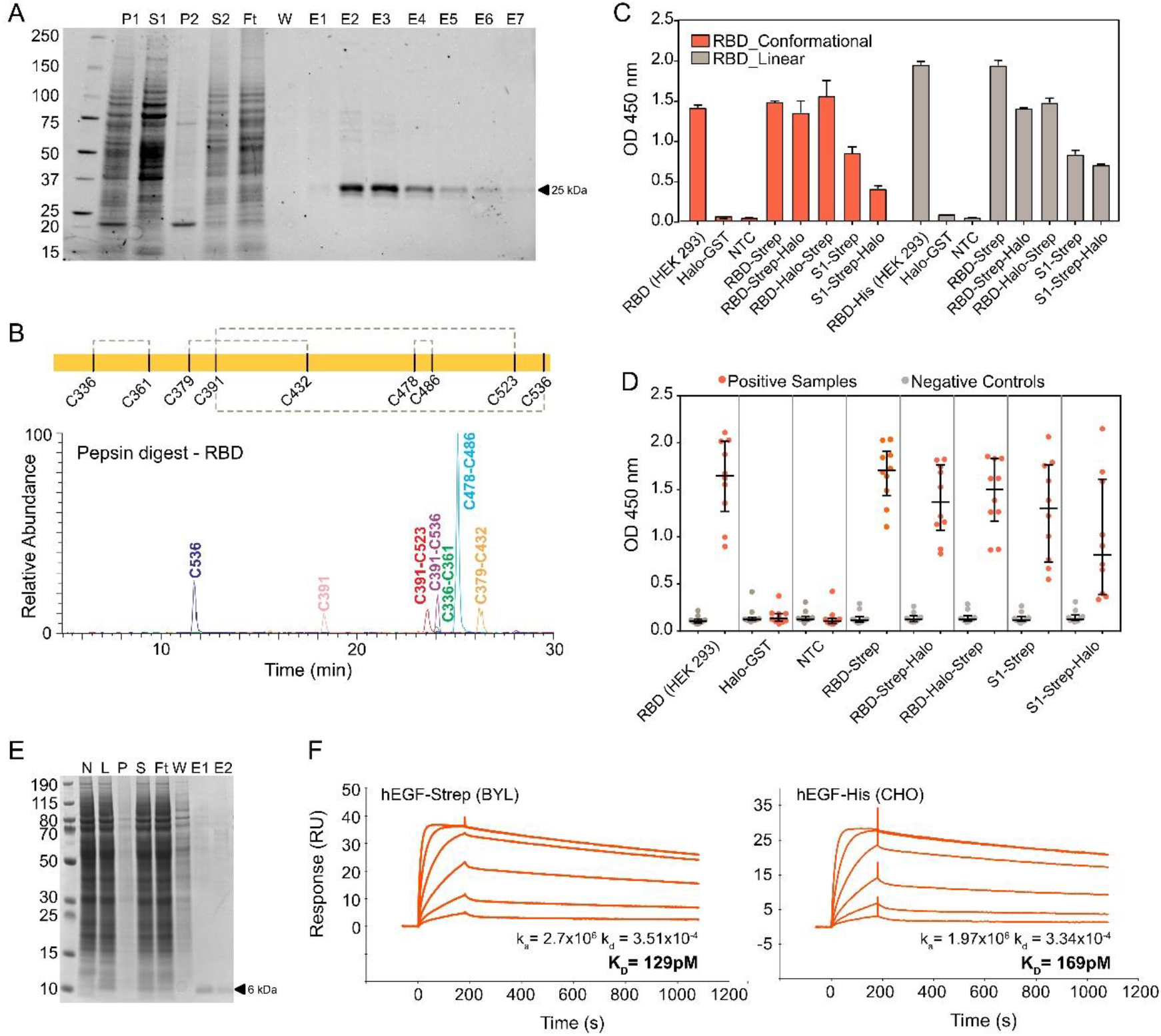
A) SDS-PAGE of RBD purification fractions. P1/P2 – insoluble fraction pellets; S1/S2 – soluble fractions; Ft – resin flow through; W – wash; E1-E7 – elution fractions. B) Disulfide bond analysis of adalimumab. C) Binding assays of SARS-CoV-2 antigens. D) Serology-based ELISA of SARS-CoV-2 antigen binding. E) SDS-PAGE of hEGF purification fractions. N – BYL non-template control; L – BYL lysate after CFPS reaction; P – insoluble fraction pellet; S – soluble fraction; Ft – resin flow through; W – wash; E1/E2 – elution fractions. F) SPR comparison of hEGF produced in either BYL or *E. coli*.

This protein is considered an important target for developing a SARS subunit vaccine. Indeed, the majority of the potent neutralising antibodies against SARS-CoV-2 target the RBD^30^, but reports differ on whether RBD alone or full S1 subunits of the spike are more potent vaccines in terms of IgG and IgA induction^31,32^.To explore this concept, we expressed both RBD and this S1 subunit from the pALiCE02 vector and with different HaloTag and Strep-II tag conformations. Antigen reactivity to two commercial antibodies was initially assessed, selected based on the recognition of either linear or conformational RBD epitopes. We observed similar reactivity of both antibodies against recombinantly expressed RBD and S1 from both BYL CFPS and HEK293 cells (Figure 6C). Importantly, the reactivity against S1 was weaker than that for RBD.

Furthermore, to demonstrate true biological relevance of proteins produced in BYL, we found clear serological reactivity of the antigen samples (Figure 6D). An adapted ELISA workflow utilizing SARS-CoV-2 patient serum antibodies showed that the RBD produced in BYL was directly comparable to that of RBD produced in HEK293 cells. Interestingly, higher reactivity was observed for COVID-19 patient samples but lower reactivity for samples collected from control subjects prior to 2019, and this separation was less signification for S1 antigens. Different C-terminal Strep-II tag and HaloTag configurations did not affect binding, suggesting correct RBD folding independent of fusion protein additions. This experiment suggests the use of RBD produced using BYL for therapeutic purposes and scaled expression for animal studies should be a priority for future research.

Finally, human epidermal growth factor (hEGF) has been extensively investigated for its potential ability to promote rapid healing of serious injuries, such as cuts, burns, and diabetic ulcers^33^. Although hEGF promises potential clinical value, the growth factor is restricted to the treatment of chronic diabetic ulcers because of its high production cost and poor stability. Mature hEGF is a complex peptide of 53 amino acids that possess three intramolecular disulfide bonds. It was chosen as a model protein to challenge the capabilities of the BYL system as an alternative expression host for bioactive hEGF^34^.

The hEGF gene with an N-terminal Strep-II tag was cloned into pALiCE02 for microsomal expression. Single-step streptavidin affinity purification was sufficient to extract soluble hEGF at high purity (Figure 6E). Binding of the purified protein to the cognate epidermal growth factor receptor (EGFR) was used as a surrogate measure of bioactivity and analysed by SPR (Figure 6F). hEGF expressed in the BYL showed comparable or improved binding kinetics compared to a commercial hEGF reference product expressed from *E. coli*, suggesting correct folding and disulfide bond formation. Prokaryotic CFPS systems require extensive engineering to enable disulfide bond formation that can be circumvented by the use of BYL microsomes.

## Discussion

The promise of cell-free systems is acceleration of the protein biotechnology industry - anything that can be done using cells could be done faster and with lower technical and infrastructural barriers to entry using CFPS. Cell-free production of protein therapeutics and drug screening targets are clear opportunities but the fields of synthetic biology, metabolic engineering, structural biology and more all stand to benefit. However, despite many reports of independently produced cell-free lysates derived from multiple organisms, and the increasing commercial availability of cell-free kits, broad uptake of the technology in academic and industrial sectors has lagged.

Major challenges still exist for CFPS in terms of batch-to-batch consistency, yield, scalability, cost effectiveness, protein folding and protein functionality. Herein, we have disclosed the results of a decade-long development path towards solving these issues in the BY-2 cell lysate that is commercialised as ALiCE^®^. Using multiple batches of the publicly available, commercial-grade lysate, functional expression of an: enzyme, monoclonal antibody, viral antigen, VLP and membrane protein was achieved, with inter-batch comparability demonstrated across protein quality metrics of glycosylation and disulfide bonding. Linear scaling of reactions for reporter proteins across microliter to milliliter volumes has also been shown, to include 1 L reactions from a recently scaled lysate production methodology. Reactions were performed in batch mode by simply combining lysate with template DNA and incubating the mixtures, albeit with specific bioreactor equipment requirements at the liter scale. Together, the results suggest this CFPS system as an end-to-end R&D platform. First-line protein engineering and lead development activities are achievable on the microscale before transitioning to gram scale production of protein material, without requiring extensive process development and optimisation.

These varied proteins were produced within 48 hour reaction timeframes from off-the-shelf lysate starting material. However, as an open system, CFPS possesses deep opportunities for protein-specific optimisation for enhanced yields and quality attributes. Reactions can be abstracted as chemoenzymatic processes instead of biological ‘black boxes’ and more extensive physicochemical interventions are permitted without consideration of cell walls or membranes e.g. modulation of physicochemical parameters of pH and temperature, or the facile addition of cofactor and chaperone molecules^35^.

Similarly, proteins produced using CFPS can be processed for downstream applications like purification and functional analysis without the necessity of transfection, selection, and expansion of clones. This circumvents confounding variables like transfection efficiencies, indicating a more flexible screening platform than cell-based methods and with particular relevance to the monoclonal antibody discovery sector. Streamlined protein variant expression also has relevance to pandemic preparedness e.g. rapid production of SARS-CoV-2 RBD mutants from emerging viral strains for diagnostic or therapeutic purposes^36^. The use of CFPS has even been posited for individualised vaccines and distributed medicines manufacturing, presenting new medical paradigms if regulation can be safely adapted to meet this potential^37–39^.

For this study, eukaryotic model proteins were deliberately chosen to challenge the capabilities of the system. We have shown that BYL can produce functional proteins as diverse as enzymes, multi-pass transmembrane GPCRs, small, soluble growth factors and immunologically-active viral antigens. Moreover, functional expression of various biosynthetic enzymes, Cas9 nuclease and the antibody M12 have previously been reported from BYL^10,40,41^. Protein function is intrinsically linked to correct folding but further molecular characterisation was warranted to prove that the microsomes of this eukaryotic CFPS system are able to produce recombinant proteins at the same quality as a cell and with batch-to-batch consistency. Essential features such as N-linked glycosylation are not possible in prokaryotic CFPS without extensive engineering but was confirmed herein with high structural homogeneity and N-glycosite occupancy. High-mannose structures typical of plant N-glycosylation were understandably prevalent, considering the BY-2 cell culture starting material. Humanized glycosylation strategies are being explored for applications where the functional relevance of N-glycosylation is more important e.g. antibody-dependent cell-mediated cytotoxicity. There is a strong field of Nicotiana spp. and BY-2 cell glycoengineering from which to draw upon^42–46^, and we have already produced BYL from cells with genetic knock-out of plant-specific xylosyl- and fucosyltransferases.

Regardless of expression host, membrane proteins are universally considered as particularly challenging targets for recombinant expression due to specific requirements for folding, PTMs and correct insertion into cell membranes or detergent micelles^23,24^. Indeed, membrane proteins are underrepresented in the Protein Data Bank, despite constituting about 30% of the human proteome. This gap is of keen interest for the pharmaceutical industry, as 50% of approved pharmaceutical molecules target membrane proteins^47^. Functional CB2 was expressed in this study using the standard format of BYL lysate, without supplementation of exogeneous lipidic platforms or detergents, and performed equally or better than the same construct produced in standard *E. coli* culture. Additional work could be required to solubilise CB2 for the various screening and structural biology workflows that are utilised in small molecule drug discovery but it is encouraging that functional starting material can be easily obtained in BYL, whereby necessary overexpression of membrane proteins has been repeatedly posited as toxic to cell-based systems. With increasing reports of functional GPCR expression in cell-free systems, it is envisaged that this knowledge gap between membrane protein genomics and useful structures with soon be bridged by CFPS^48^.

Beyond proteins as target molecules, cell-free is also proving useful for prototyping of biosynthetic enzyme pathways for the production of high-value chemicals ^40,49^. Prototyping is enabled by throughput and whilst microfluidics show increasing potential for CFPS workflows, the equipment is broadly inaccessible in standardized laboratory setups. More universal multiwell-based screening platforms require large volumes of high-yielding lysate with dependably reproducible quality in order to effectively sample the immense experimental design space when reengineering multistep biosynthetic pathways. Consequently, our work in scaling of lysate production and optimization of product consistency promotes the application of BY-2 lysate and CFPS as a whole, to accelerate research and development pipelines for faster time-to-market turnaround of novel protein therapeutics, diagnostics and even high value small molecules.

## Materials and Methods

### Tobacco BY-2 cell culture and lysate production

Tobacco cells *(Nicotiana tabacum* L. cv. Bright Yellow - 2; BY-2) were routinely cultivated in Murashige-Skoog liquid medium (MS) at 1 L scale maintaining the exponential growth phase of the cells with a maximum packed cell volume of 40% at 26°C in the dark^50,51^. Production cultures were inoculated 3.5% in MS medium supplemented with 2% (w/v) sucrose and Pluronic L-61 antifoam and cultured for 72 h at 26°C in darkness before lysate bioprocessing. Standard lysates were prepared from liter scale cultures. For the novel, scaled lysate bioprocessing methodology, larger culture volumes were obtained from cell culture seed trains generated in commercial bioreactors. General BY-2 lysate preparation has previously been described^9,10^. Scaled lysate production was performed using a novel, commercially sensitive method.

### Gene cloning and DNA template preparation

Genes encoding proteins of interest were purchased from gene synthesis providers (Integrated DNA Technologies, Inc.; Twist Bioscience) and transferred into pALiCE01 and pALiCE02 expression vectors (LenioBio GmbH) using the Gibson DNA assembly method^52^. Vectors provided proteins of interest with N- or C-terminal Strep-II tags to enable facile downstream purification, except in the case of adalimumab where tag-less constructs were used. DNA fragments were amplified with 15-20 bp of homology regions in primers using Phusion R High Fidelity DNA polymerase (New England Biolabs; NEB) and homology based assemblies were performed using NEBuilder® HiFi DNA Assembly Master Mix (NEB) following manufacturer guidelines and transformed into *E. coli* DH5-alpha cells. Correct clones were determined by sequencing and cultured for subsequent plasmid preparation using NucleoBond Xtra Midi/Maxi kits (Macherey-Nagel GmbH & Co. KG). For scaled CFPS reactions, milligram quantity production of template DNA was contracted to PlasmidFactory GmbH & Co. KG..

### Cell-free protein synthesis reactions using BY-2 lysate

Generally, CFPS reactions were performed according to the ALiCE® instruction manual and published protocols^52^. Briefly, relevant volumes of BYL product were thawed in a room temperature water bath and template DNA added to final concentration of 5 nM (excepting the heteromeric adalimumab, whereby 5 nM of each heavy and light chain plasmids were used for a total DNA concentration of 10 nM). ‘Non-template controls’ lacking DNA were always included to provide lysate background samples for downstream assays. Incubation temperature and runtime of 25 °C and 48 hours were used, irrespective of reaction volumes.

Microscale CFPS reactions of 50 and 100 μL were performed in microtiter plates using a shaking speed of 700 rpm. Additionally, Duetz system microtiter plate covers were used to limit uneven evaporation and edge effects (EnzyScreen BV). At the millilitre scale, reactions were optimised using Erlenmeyer type glass flasks. CFPS reactions of 10 and 100 mL were performed using sterilised 250 and 5000 mL flasks, respectively, with a shaking speed of 105 rpm. Finally, the 1 L scale reaction was performed using a CELL-tainer® CT20 rocking motion bioreactor with single-use bag.

### Reporter protein quantification

Expression of reporter proteins was quantified directly from lysate samples after CFPS reactions. For eYFP, lysate samples were diluted 1:10 with 20 mM MOPS buffer at pH 7.2 in microtiter plates. A calibration curve of eYFP standard was similarly prepared and the plate scanned for fluorescence signal with excitation and emission wavelengths of 485/20 and 528/20 nm, respectively, using an Infinite M1000 device (Tecan Group Ltd.). Quantification was achieved by subtracting raw fluorescence signals for non-template control from the values acquired for test samples and interpolating the adjusted value against the calibration curve.

GOx expression was quantified through a horseradish peroxidase (HRP)-coupled activity assay. Microsomal proteins were released by treatment of the lysate samples with 0.5% dodecylmaltoside (DDM) detergent for at least 15 minutes at room temperature. Samples were diluted 1:2500 in an assay buffer comprising 0.33 M glucose, 0.67 mM ABTS and 1.67 U/ml HRP (Sigma Aldrich) in 0.1 M potassium phosphate buffer at pH 6 in transparent 96 well plates. Absorbance over time at 420 nm was measured using the Infinite M1000 plate reader every 15 seconds over a period of 15 minutes. The linear absorbance increase over time was used to calculate sample GOx activity.

### SDS-PAGE and Western blot analyses

Precast 4–15% Mini-PROTEAN™ TGX Stain-Free™ (Bio-Rad Laboratories, Inc.) or NuPAGE 4-12% Bis-Tris (Thermo Fisher Scientific) were used for SDS-PAGE. Stain-free gels were visualised under UV light but were otherwise stained with Coomassie brilliant blue R-250. PageRuler™ Plus Prestained Protein Ladder 10-250 kDa (Thermo Fisher Scientific) was used as molecular weight marker.

Proteins were transferred from gels to PVDF membranes using a Trans-Blot® Turbo™ Transfer System and associated accessories (Bio-Rad Laboratories, Inc.). Membranes were probed with anti-Strep-II tag and anti-CB2 antibodies according to manufacturer instructions.

### Production and Transmission Electron Microscopy of Virus-like Particles

The human hepatitis B virus core antigen (HBc; serotype adw) gene was cloned into the pALiCE01 vector, generating the pALiCE01-HBc plasmid. Microscale BYL reactions of 100 μL reactions were performed using this template DNA. Resulting lysate samples were processed by centrifuging at 16,000x *g* for 10 minutes, to clarify the non-soluble fraction. Supernatant containing VLPs were collected and the non-soluble pellet resuspended in phosphate buffered saline (PBS). Each fraction was analysed by SDS-PAGE.

For negative staining electron microscopy (EM), formvar/carbon coated 400 mesh copper grids were exposed to a glow discharge in air for 20 seconds before applying 10 μL of the VLP suspensions. Grids were incubated at room temperature for 2 minutes and negative staining was performed with 1% phosphotungstic acid, pH 7.2, for 1 minute. The specimens were examined in a JEM-1400Flash Electron Microscope equipped with a Matataki Flash 2k x 2k camera (JEOL, Ltd.).

### CB2 expression and functional assay

The full-length human cannabinoid type 2 (CB2) receptor was expressed in *Escherichia coli* as a fusion with the N-terminal maltose-binding protein (MBP), Strep-II tag and C-terminal deca-histidine affinity tags (MBP-StrepII-CB2-His_10_), and human embryonic kidney (HEK) Expi239 cells without the MBP module (StrepII-CB2-His_10_). Expression and purification from cell membranes was performed as previously described^27,53^. CB2 in complex with E. coli membranes was also isolated. The MBP-StrepII-CB2-His_10_ construct was cloned into pALiCE01 and pALiCE02 vectors for subsequent microscale expression in 50 μL reactions. Resulting BYL samples and purified or membrane-bound CB2 proteins were analysed by SDS-PAGE and Western blotting with an anti-CB2 and anti-Strep-II antibodies.

Activation of G proteins by the recombinant CB2 was performed according to a previously reported protocol^28^. Briefly, BYL samples after CB2 expression or purified CB2 protein in Façade-TEG/ CHS micelles were dispensed into pre-siliconized glass tubes containing 10 mM MOPS supplemented with 0.1% (w/v) BSA and 10 μM CP-55,940 (Cayman Chemical). Upon addition of a mixture of G_αi1_ (100 nM) and G_β1γ2_ (500 nM), the tubes were incubated on ice for 30 min. Reactions were completed to a total volume of 50 μL with 50 mM MOPS buffer at pH 7.5, 1 mM EDTA, 3 mM MgCl_2_, 4 μM GDP, 0.3% w/v BSA, 100 mM NaCl, 1 mM DTT and an appropriate amount of ^35^S-γ-GTP, with tubes transferred rapidly to a 30 °C water bath. Incubation continued for 20 minutes and reactions terminated by addition of 2 mL ice-cold stop solution TNMg (20 mM Tris–HCl pH 8.0, 100 mM NaCl, 25 mM MgCl2). The reaction was rapidly filtered through 0.45 μm nitrocellulose filters. Filters were washed with 4 × 2 mL of cold TNMg buffer, dried, placed into scintillation vials and counted upon addition of ScintiSafe Econo F scintillation liquid (Fisher).

### Purification of microsomal proteins from BYL lysate

Medium scale reactions of 10 mL were used to express human epidermal growth factor (hEGF), adalimumab, GOx and SARS-CoV-2 receptor binding domain (RBD) from pALiCE02 vectors. Following the CFPS reaction, lysates were centrifuged at 16,000x *g* for 20 minutes at 4 °C to pellet the microsomal fraction. Microsome pellets were resuspended in 0.5% DDM in PBS to the original volume of the reaction and incubated for at least 15 minutes at room temperature to disrupt the microsome and release the soluble protein. Suspensions were then clarified by a repeated centrifugation step an the supernatant progressed for purification.

nProtein A Sepharose 4 Fast Flow resin (Cytiva) was used for affinity purification of adalimumab whilst Strep-Tactin XT 4Flow resin (IBA Lifesciences) was used for Strep-II tagged-tagged proteins. Purifications were performed manually according to manufacturer instructions and resulting elutions were concentrated and buffer exchanged to PBS using Amicon Ultra Centrifugal Filter Units (Merck). Purified, concentrated protein samples were calculated by comparing absorbance measurements against theoretical molar extinction coefficients for specific proteins.

### Surface plasmon resonance analyses

SPR spectroscopy was conducted on a Biacore T200 instrument (Cytiva). HBS-EP+ at pH7.4 (Biacore/Cytiva) was used as the running buffer, with measurements were conducted at 25°C and at a flow rate of 30μL/min. For capture of antibodies and Fc-Fusions a CM5-S-Series sensor chip (Cytiva) was functionalized with 2500RU recombinant Protein A (Sigma) using EDC/NHS Coupling Kit (Biacore/Cytiva BR-1000-50) according to the manufacturers suggestion. Flow cell one (Fc 1) was activated and deactivated to be used as a reference for blank subtraction. Between the measurement cycles, the surface was regenerated by pulsing for 45 sec with the recommended regeneration buffer (10 mM glycine-HCl pH 2.1 supplied within the Human Fab Capture Kit). Buffer injections were used for double referencing.

Kinetic parameters were determined for the interaction of BYL-derived adalimumab batches as well as commercial Humira® (antibodies-online) with the corresponding antigen tumor necrosis factor (TNFα; Abcam plc), as well as with the human FcR I (CD64; R&D Systems) receptor. Antibodies were captured by injecting an appropriate dilution of each respective molecule in running buffer for 180 seconds, followed by an injection of either TNFα or CD64 for 180 seconds followed by a 420 second dissociation phase and a 45 second regeneration step using 30mM HCl. The cycle was repeated with eight serial twofold dilutions of the respective ligands, starting at 60nM for hTNFalpa and 30nM for hCD64. Affinity constants were determined through fitting the resulting sensograms with the BiacoreEval 3.0 software after performing double-referencing using the Biocore T200 evaluation software. A 1:1 binding model was used.

A similar approach was used to quantify the interaction between BYL-derived hEGF and a standard from E. coli (R&D Systems) with recombinant EGFR Fc-fusion (R&D Systems). EGFR was captured by injecting an appropriate dilution of the molecule in running buffer for followed by injection of the hEGF samples following the parameters outlined above. The cycle was repeated with five serial twofold dilutions of the ligands, starting at 41,6nM. Affinity constants were determined as above.

### Deglycosylation Assay, SDS-PAGE and Immunoblot

Deglycosylation with either EndoH or PNGaseF (NEB) was performed according to the manufacturer’s instructions. Aliquots of purified proteins were treated with either EndoH or PNGaseF, followed by SDS-PAGE analysis.

### N-glycan and disulfide bond analyses via mass spectrometry

For N-glycopeptide analyses, purified protein samples were prepared by S-alkylation with iodoacetamide and digestion with either with LysC/GluC (Promega) [RBD] or, in the case of adalimumab, trypsin (Promega). Samples were loaded on a nanoEase C18 column (nanoEase M/Z HSS T3 Column, 100Å, 1.8 μm, 300 μm X 150 mm, Waters) using 0.1 % formic acid as the aqueous solvent. A gradient from 1% B (B: 80% Acetonitrile, 0.1% FA) to 40% B in 50 min was applied, followed by a 10 min gradient from 40% B to 95% B that facilitates elution of large peptides, at a flow rate of 6 μL/min. Detection was performed with an Orbitap MS (Exploris 480, Thermo Fisher Scientific) equipped with the standard H-ESI source in positive ion, DDA mode (= switching to MSMS mode for eluting peaks). MS-scans were recorded (range: 350-1200 Da) and the 20 highest peaks were selected for fragmentation. Instrument calibration was performed using Pierce FlexMix Calibration Solution (Thermo Fisher Scientific). The possible glycopeptides were identified as sets of peaks consisting of the peptide moiety and the attached N-glycan varying in the number of HexNAc units, hexose, deoxyhexose and pentose residues. The theoretical masses of these glycopeptides were determined with a spread sheet using the monoisotopic masses for amino acids and monosaccharides. Manual glycopeptide searches were made using Freestyle 1.8 (Thermo Fisher Scientific). For the quantification of the different glycoforms the peak areas of EICs (Extracted Ion Chromatograms) of the first four isotopic peaks were summed, using the quantification software Skyline.

Disulfide bond analysis workflows were conducted similarly. Protein samples were S-alkylated and processed with pepsin or, in the case of adalimumab, AccuMAP™ Low pH Protein Digestion Kit (Promega) according to the manufactures protocol. The digested samples were loaded on a nanoEase C18 column with separation and detection and separated using identical parameters as above. The files were searched against databases containing the primary sequences for the full-length proteins.

### SARS-CoV-2 RBD Binding and Serological Assay

Anti-RBD antibodies were procured to recognise either linear or conformation epitopes (MAB10540 and MAB105802, respectively; R&D Systems). Serum samples were collected either from COVID-19 patients with a positive PCR test in 2020 or prior to 2019 as negative controls. HaloTag-GST was used as a negative control antigen (Promega).

Pure protein samples were coated to microtiter plates overnight with 4°C incubation at a concentration of 2 μg/mL. After washing with 0.2% Tween-20 in PBS (PBST) and blocking with PBST-1% BSA, antibodies at 1:10000 dilution or serum samples at 1:200 dilution were introduced and incubated at room temperature for 1 hour. After further 0.2 % PBST washing, HRP-conjugated anti-mouse IgG or anti-human IgG detection antibodies (Sigma) were added at 1:10000 dilution with a final room temperature incubation for 1 hour. Plates were then developed with 3,3’,5,5’-tetramethylbenzidine substrate and absorbance readings at 450 nm measured using an Envision plate reader.

## Acknowledgements

This project has received funding from the European Union’s Horizon 2020 research and innovation program, under grant agreement No 881025 ‘PEPPER’, and from the German BMBF Biooekonomie No 0312B083OB ‘ThinkBig’ project. We thank our colleagues at LenioBio for their support and expertise during the performance of this research. We greatly appreciate the contribution of Dr. Jan W.M. van Lent and the Wageningen Electron Microscopy Center (WEMC), Wageningen, Netherlands for the electron microscopy pictures and wish him a very happy retirement. Thanks also to Clemens Grünwald-Gruber and the Mass Spectrometry core facility of BOKU, Austria for their work in mass spectrometry analyses.

